# Activated Expression of Master Regulator MYB31 and of Capsaicinoid Biosynthesis Genes Results in Capsaicinoid Biosynthesis and Accumulation in the Pericarp of the Extremely Pungent *Capsicum chinense*

**DOI:** 10.1101/2020.11.05.369454

**Authors:** Binmei Sun, Zubing Huang, Juntao Wang, Jianlang Wei, Wen Cai, Yuan Yuan, Shuangling Zhang, Jiali Song, Bihao Cao, Changming Chen, Panrong Cao, Guoju Chen, Jianjun Lei, Zhangsheng Zhu

**Affiliations:** Key Laboratory of Biology and Germplasm Improvement of Horticultural Crops in South China, Ministry of Agriculture, College of Horticulture, South China Agricultural University, Guangzhou, 510642 China; Peking University-Southern University of Science and Technology Joint Institute of Plant and Food Sciences, Department of Biology, Southern University of Science and Technology, Shenzhen 518055, China; Henry School of Agricultural Science and Engineering, Shaoguan University, Guangdong 512005, China

**Keywords:** Hot pepper, capsaicinoids, pericarp, capsaicinoid biosynthesis genes, master regulator MYB31

## Abstract

Capsaicinoids confer pungency in *Capsicum* fruits, and the capsaicinoid content varies greatly among the five domesticated *Capsicum* species. Although it is generally recognized that capsaicinoid biosynthesis occurs exclusively in the placenta, few studies have focused on capsaicinoid biosynthesis gene (CBG) expression in the pericarp. Therefore, the transcriptional regulation mechanisms of capsaicinoid biosynthesis in the pericarp remain elusive. Here, the capsaicinoid contents of 32 accessions from five domesticated *Capsicum* species were analyzed. The results showed that the capsaicinoid contents of *C. chinense* accessions are significantly higher than those of the other four *Capsicum* species due to the increased accumulation of capsaicinoids, especially in the pericarp. Compared to that in accessions with low pericarp capsaicinoid content, the expression of the master regulator *MYB31* is significantly upregulated in the pericarp in *C. chinense* accessions, which leads to high levels of CBG expression. Moreover, in fruits of the extremely pungent ‘Trinidad Moruga Scorpion’ (*C. chinense*) and low-pungency ‘59’ inbred line (*C. annuum*) at different developmental stages, the capsaicinoid accumulation patterns were consistent with the *MYB31* and CBG expression levels in the pericarp. Taken together, our results provide novel insights into the molecular mechanism arising from the expression of a master regulator in the pericarp that results in exceedingly hot peppers. The genetic resources identified in this study could be used as genetic resources for the genetic improvement of pepper pungency.

## Introduction

Pepper (*Capsicum*) is an economically important horticultural plant that is used worldwide as a vegetable and food additive owing to its unique pigment, flavor, and aroma (Pino et al., 2007). Furthermore, pepper is also grown for chemical and medicinal industries because it is extraordinarily rich in secondary metabolites (Wahyuni et al., 2011). *Capsicum* fruits contain high levels of vitamins A and C, carotenoids, and flavonoids (Ananthan et al., 2018; Liu et al., 2020; Sweat et al., 2016; Wahyuni et al., 2011). Hot pepper fruits also produce the species-specific compounds capsaicinoids, which confer a hot or pungent flavor (Zhu, Sun, Cai, et al., 2019). Capsaicinoids are a group of alkaloids and serve as deterrents against fungi, bacteria and mammals, protecting the fruit from damage (Tewksbury et al., 2008). In addition, capsaicinoids are valuable compounds that are widely used in food additives, chemicals, pharmaceuticals and police equipment (Zhu, Sun, Cai, et al., 2019). The most common capsaicinoids are capsaicin (CAP) and dihydrocapsaicin (DHCAP), accounting for approximately 91% of the capsaicinoids in *Capsicum* species (Duelund & Mouritsen, 2017; Wahyuni et al., 2011). *Capsicum* has approximately 35 species and is native to Central America (Carrizo García et al., 2016). Among them, *C. annuum*, *C. baccatum*, *C. chinense*, *C. frutescens*, and *C. pubescens* are widely cultivated in different parts of the world. The capsaicinoid content fluctuates greatly among these five domesticated *Capsicum* species (Sweat et al., 2016; Wahyuni et al., 2011; Zhu, Sun, Cai, et al., 2019). Genetics is considered the most important factor that determines the pungency level of peppers (Wahyuni et al., 2011; Zhu, Sun, Cai, et al., 2019). Currently, all the hottest identified pepper varieties (such as ‘Habanero’, ‘Bhut Jolokia’ and ‘Trinidad Moruga Scorpion’) are *C. chinense* species (Bosland et al., 2012; Bosland & Baral, 2007; Canto-Flick et al., 2008; Duelund & Mouritsen, 2017). In addition, they are also influenced by endogenous and exogenous factors, such as fruit developmental stages, phytohormones and growth conditions (N. Jeeatid et al., 2018; Nakarin Jeeatid et al., 2017; Naves et al., 2019; Phimchan et al., 2012; B. Sun et al., 2019; Binmei Sun et al., 2019). Light, temperature, water and nutrients influence capsaicinoid biosynthesis, and these cultivation conditions govern capsaicinoid biosynthesis in a genotype-dependent manner (N. Jeeatid et al., 2018; Nakarin Jeeatid et al., 2017; Naves et al., 2019). Therefore, there is no consensus conclusion regarding the effects of environmental factors on capsaicinoid accumulation (Naves et al., 2019). Capsaicinoid content is greatly influenced by growth conditions, but the officially reported hottest *Capsicum* cultivars have not been systematically compared under the same growth conditions. In addition, for some cultivars, such as ‘Shuanla’ (*C. chinense*), which is the hottest pepper in China (Deng et al., 2009), its world pungency ranking remains unknown. The understanding of capsaicinoid contents in different genotypes is a precondition for the genetic improvement of the pungency level in *Capsicum*.

The capsaicinoid biosynthesis pathway has been almost completely determined and is composed of the phenylpropanoid and branched-chain fatty acid pathways (Kim et al., 2014). Many genes, such as *Kas*, *AMT*, *KR1*, *ACL*, *FatA* and *AT3* (also known as *Pun1*), have been reported to play crucial roles in capsaicinoid biosynthesis, and their expression levels are significantly correlated with capsaicinoid content (Aluru et al., 2003; Curry et al., 1999; Del Rosario Abraham-Juárez et al., 2008; Qin et al., 2014; Stewart et al., 2005). Capsaicinoid levels and CBG transcription levels are highly dynamic during fruit development (Kim et al., 2014; Zhu, Sun, Cai, et al., 2019). Capsaicinoids begin to accumulate at the early developmental stage of fruit and peak at approximately 45 days postanthesis (DPA) (Barbero et al., 2014; Fayos et al., 2019; Zhu, Sun, Cai, et al., 2019), but the precise accumulation pattern is genotype-dependent. Accordingly, CBGs are also expressed during the development of the pepper placenta and facilitate capsaicinoid synthesis in the placenta throughout this period (Kim et al., 2014; Zhu, Sun, Cai, et al., 2019). In addition, *MYB31* has been reported to be a master regulator of capsaicinoid biosynthesis; it directly binds to the CBG promoters to regulate capsaicinoid biosynthesis (Arce-Rodríguez & Ochoa-Alejo, 2017; Zhu, Sun, Cai, et al., 2019). The loss of function of *MYB31* in the accession ‘YCM334’ (*C. annuum*) leads to significantly downregulated CBG expression, which finally results in the fruit losing pungency(Han et al., 2019). Generally, capsaicinoid biosynthesis occurs exclusively in the placenta, and CBGs are specifically expressed in this tissue (Fujiwake et al., 1980; Suzuki et al., 1980). Recently, a published study showed that CBGs are also expressed in the pericarp of the cultivar ‘Trinidad Moruga Scorpion Yellow’ (*C. chinense*), which suggests capsaicinoid biosynthesis in the pericarp (Tanaka et al., 2017), but the underlying transcriptional regulation mechanisms for capsaicinoid accumulation have yet to be elucidated. In addition, whether the CBGs are consistently upregulated in the pericarp of other extremely pungent cultivars and whether *MYB31* executes its functions to control CBG expression in the pericarp is still unknown. Therefore, the relationship of *MYB31* and CBG expression to capsaicinoid content in five domesticated *Capsicum* pericarps needs to be addressed.

In this study, the capsaicinoid contents of 32 accessions, including some of the officially reported hottest accessions from five *Capsicum* species, were determined. By analyzing the capsaicinoid contents in the placenta, pericarp and whole fruit of the 32 accessions, we revealed that capsaicinoids are highly accumulated in accessions of the *C. chinense* pericarp. In addition, the increased pungency in the accessions of *C. chinense* pericarp was caused by the upregulation of *MYB31* and CBGs. In addition, the capsaicinoid accumulation pattern and *MYB31* and CBG expression levels in extremely pungent ‘Trinidad Moruga Scorpion’ and low-pungency ‘59’ inbred line fruit at different developmental stages in the pericarp were also characterized. Our results provide novel insight into the transcriptional regulation of capsaicinoids in the pericarp and valuable references for breeders improving the pungency of peppers.

## Materials and methods

### Chemical reagents

CAP and DHCAP standards were obtained from Sigma-Aldrich (Sigma, USA). Ultrapure water was prepared by a Barnstead™ GenPure™ Pro (Thermo Fisher, USA). HPLC-grade methanol, acetonitrile, ethanol, and tetrahydrofuran were procured from Merck (Merck, USA). The solvents were filtered through a 0.22-μm nylon membrane filter (Millipore, USA).

### Plant Materials

Pepper seeds were planted in a greenhouse to grow the seedlings. The pepper seedlings were transplanted to a plastic greenhouse in an experimental field at South China Agricultural University (Guangzhou, China). The pepper plants were grown under normal conditions, and the fruits began to set in early May. The fruits were harvested at the indicated time point. For pre-extraction processing, the samples of fresh pepper pericarp, placenta and whole fruit were cut into small pieces and dried using a FreeZone freeze-dryer (Labconco, USA) until a constant weight was achieved.

### Extraction of capsaicinoids

The freeze-dried pepper fruits were ground into a fine powder, and 0.1 g of the samples was extracted with 10 ml methanol:tetrahydrofuran (1:1) based on a previous method with minimal modifications (Zhu, Sun, Cai, et al., 2019). Briefly, the extractions were performed in a Branson 8510 ultrasonic bath (320 W, 40 kHz) (Branson, USA) with a nominal transducer output amplitude of 40% at 40°C for an extraction period of 60 min to ensure that the capsaicinoids were completely extracted from the samples. For the HPLC analysis, 10 μl of CAP and DHCAP standard compound solutions (Sigma-Aldrich, USA) or extracted capsaicinoid samples were injected into an XSelect HSS C-18 SB column (4.6◻×◻250◻mm, 5◻μm, Waters Technologies). The samples were eluted at a flow rate of 1 ml/min with 20% water (A) and 80% methanol (B) as mobile phases at 30 °C on a Waters Alliance 2489 separation module (Waters, USA). Capsaicinoid detection was performed with a 2489 UV/visible detector at 280◻nm (Waters, USA). The total capsaicinoids were represented as (CAP + DHCAP)/0.9 according to previous studies (Bosland & Baral, 2007; Wahyuni et al., 2011; Zhu, Sun, Cai, et al., 2019). Finally, the CAP and DHCAP contents were converted to SHU as described previously (Collins et al., 1995).

### RNA extraction and quantitative reverse transcription PCR (qRT-PCR) analysis

The fruit placenta and pericarp were sampled and immediately frozen in liquid nitrogen. RNA was extracted using the HiPure Plant RNA Mini Kit according to the manufacturer’s protocols (Magen, China). Reverse transcription was performed using HiScript III RT SuperMix for qPCR (+gDNA wiper) (Vazyme Biotech, China). qRT-PCR analyses were performed using ChamQ SYBR qPCR Master Mix (Vazyme Biotech, China) on a CFX384 Touch (Bio-Rad, USA). The PCR conditions were as follows: 5 min at 95°C followed by 40 cycles of 10 s at 94°C, 10 s at 55°C and 20 s at 72°C. Housekeeping genes were used for gene expression analysis. The sequences of the primers used in this study are listed in Supplementary Table 1.

### Data analysis

Data are reported as the mean ± SD. Each analysis was performed in triplicate. Data were analyzed by one-ANOVA followed by a multiple comparisons test using SPASS 20 for Windows.

## Results and discussion

### Assessment of capsaicinoid content in 32 accessions from five *Capsicum* species

A total of 32 accessions (13 *C. chinense*, 9 *C. annuum*, 4 *C. frutescens*, 4 *C. baccatum*, and 2 *C. pubescens*), including the previously reported hottest cultivars (sample 1, ‘Trinidad Moruga Scorpion’ from Trinidad; sample 4, ‘Carolina Reaper’ from America; sample 9, ‘Shuanla’ from China; and ‘Bhut Jolokia’ from India) were assessed. The results showed that the content of capsaicinoids varied significantly among the 32 accessions (Fig. 2). In the placenta, CAP (6.6 mg/g to 101.4 mg/g), DHCAP (2.6 mg/g to 58.6 mg/g) and total capsaicinoid (10.3 mg/g to 165.7 mg/g) levels varied greatly among the 32 genotypes (Fig. 2A), but they were consistently higher in the accessions of *C. chinense* than in the accessions of the other four species (Fig. 2). In particular, unexpectedly high levels of capsaicinoids were detected in the *C. chinense* pericarp (Fig. 2B). In contrast, capsaicinoids were rarely found in the pericarp of the other four species’ accessions (Fig. 2B). In the *C. chinense* pericarp, the total capsaicinoid content ranged from 36.2 mg/g to 138.0 mg/g; the total capsaicinoid content ranged only from 1.38 mg/g to 7.74 mg/g in the other four *Capsicum* species (Fig. 2B). Because both the placenta and pericarp accumulate abundant capsaicinoids, the total capsaicinoid levels in the accessions of whole *C. chinense* fruit ranged from 32.4 mg/g~118.5 mg/g, which is markedly higher than the levels of 1.86 mg/g ~11.3 mg/g detected in the other four species (Fig. 2C). Notably, the highest total capsaicinoid content was observed in ‘Carolina Reaper’ (sample 4), in which the capsaicinoid content was 100-fold higher than that of the lowest, the ‘750’ inbred line (sample 19) (Fig. 2B). These findings are consistent with previous reports that ‘Carolina Reaper’ is the hottest pepper in the world (Duelund & Mouritsen, 2017). Regarding the hottest Chinese cultivar, ‘Shuanla’ (sample 9), the contents of total capsaicinoids in the placenta, pericarp and whole fruit were 133.1 mg/g, 100.8 mg/g and 68.1 mg/g, respectively. ‘Shuanla’ was less pungent than ‘Carolina Reaper’, and the total capsaicinoid contents in the placenta, pericarp and whole fruit of ‘Shuanla’ accounted for 80%, 73% and 57% of those of ‘Carolina Reaper’, respectively. In accessions of *C. chinense* species, the capsaicinoid content was at a similar level between the placenta and pericarp (Fig. 2A and 2B), and the ratio of the total capsaicinoid content in the placenta to that in the pericarp ranged from 1.02 to 2.71 (Supplementary Fig. 3). In contrast, the total capsaicinoid content in the placenta was significantly higher than that in the pericarp in the other four species (Supplementary Fig. 3), and the placenta/pericarp ratio ranged from 4.72 to 26.5. Similarly, we also found that the capsaicinoid contents of the placenta, pericarp and whole fruit in *C. chinense* accessions were relatively similar, while the contents in the placenta were significantly higher than those in the pericarp and the whole fruit in the other four *Capsicum* species (Supplementary Fig. 3). In some accessions, such as sample 8 (‘Chocolate Trinidad Scorpion’), a high level of capsaicinoids was found in the pericarp (125 mg/g) and placenta (151. 9 mg/g) but only 54.9 mg/g was found in the whole fruit because the fruit produces many seeds.

**Fig. 1.**
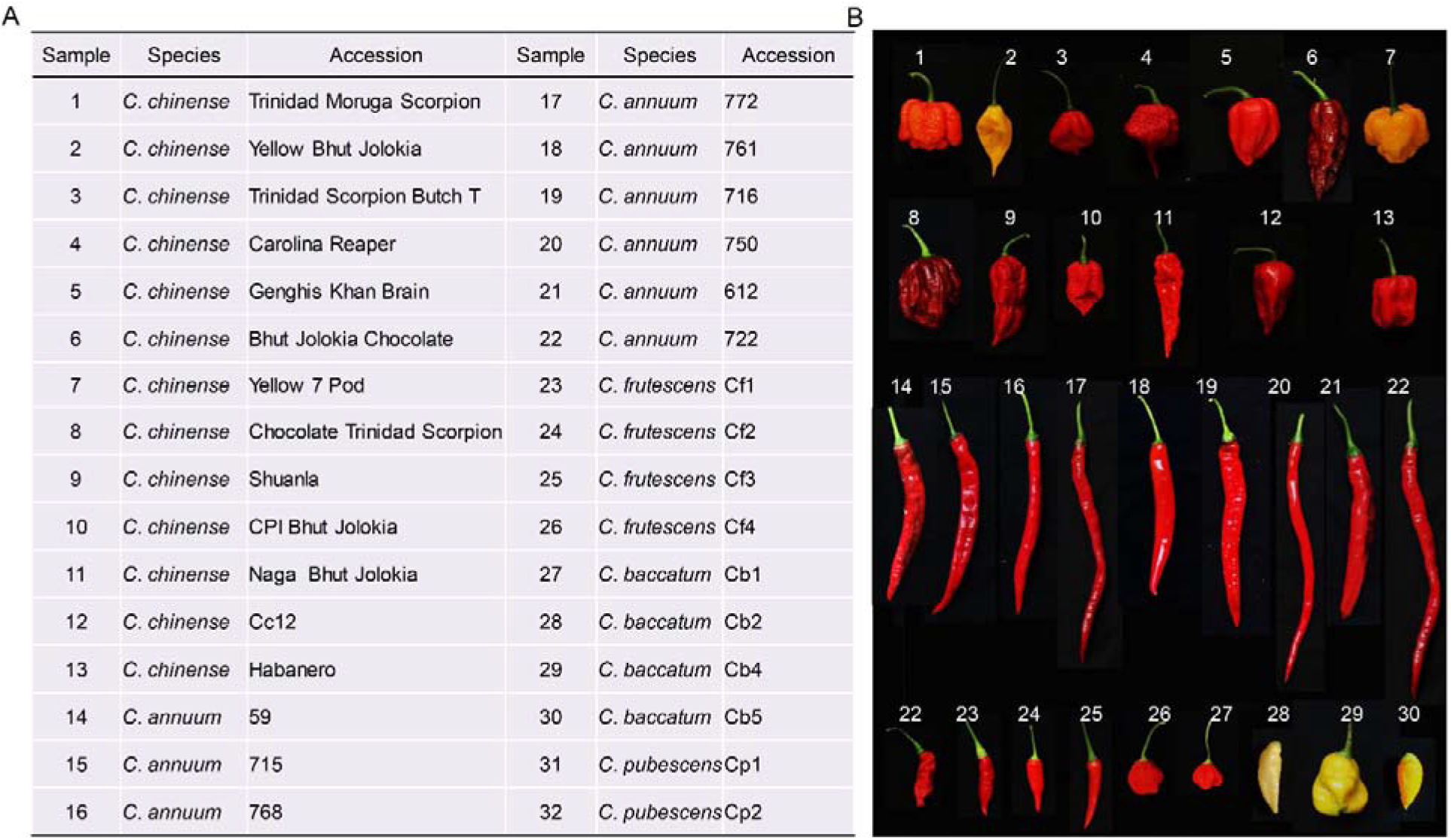
Details of the pepper accessions used for capsaicinoid analysis. (A) Pepper accessions from five *Capsicum* species. (B) The fruits of some pepper accessions exhibited full maturity at the 45 DPA developmental stage.

**Fig. 2.**
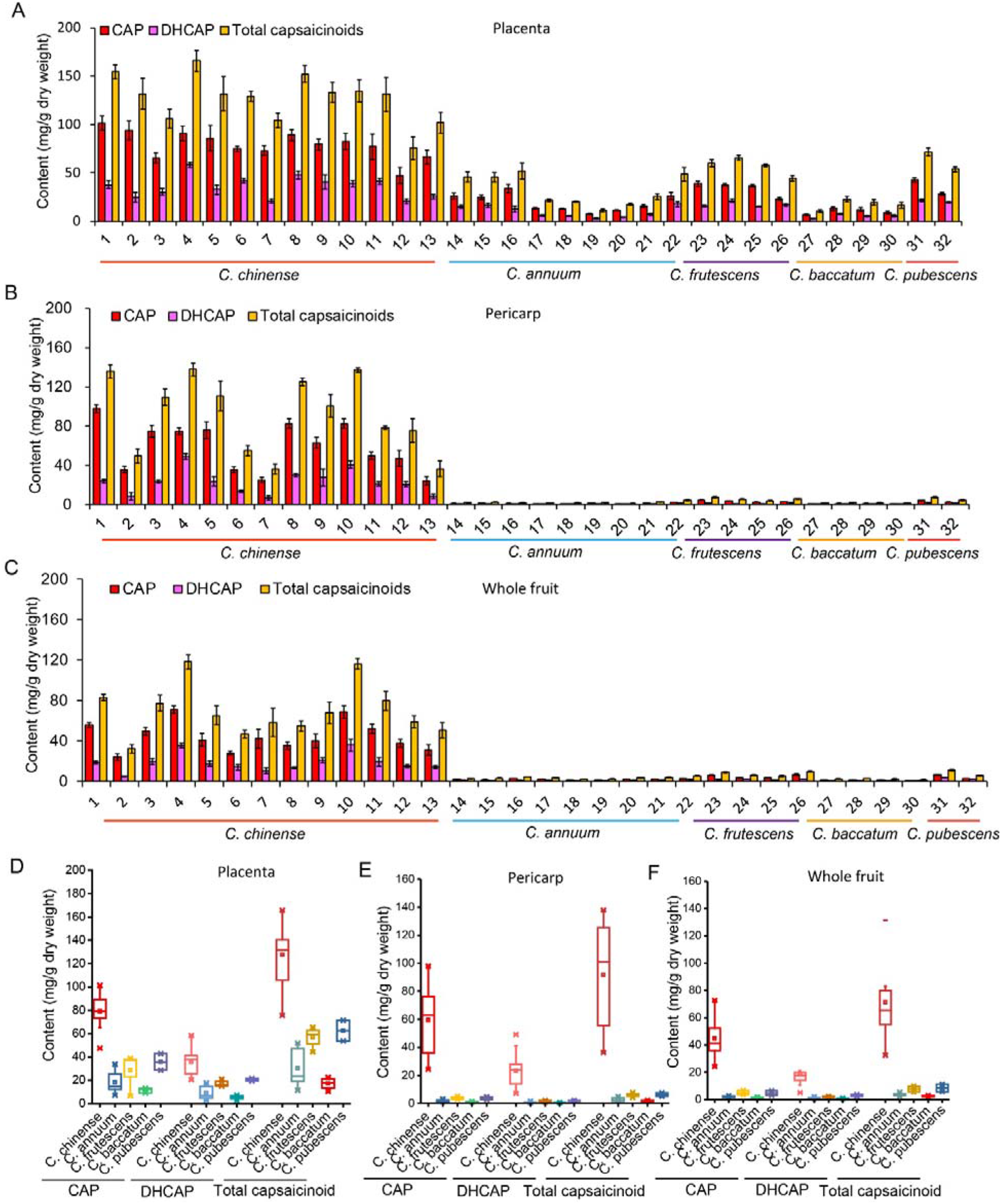
The capsaicinoid accumulation patterns in pepper fruits are species-dependent. (A-C) CAP, DHCAP and total capsaicinoid content in the pepper placenta, pericarp and whole fruit at the 45 DPA fruit development stage. Data are presented as the mean ± SD (n=3). (D-G) The box plot shows the CAP, DHCAP and total capsaicinoid content in the placenta, pericarp and whole fruit based on the analysis of the data obtained from (A-C).

To further compare the capsaicinoid content among the investigated *Capsicum* species, accessions from the same species were assigned to the same panel. The average total capsaicinoid level ranged from 17.2 mg/g (*C. baccatum*) to 127.5 mg/g (*C. chinense*) in placental tissues, and the highest levels were observed in accessions of *C. chinense* (Fig. 2D-F). Genotype (i.e., species) does seem to be a key factor determining the capsaicinoid content because the capsaicinoid levels in accessions of *C. chinense* were consistently higher than those in accessions of the other four *Capsicum* species. Previous studies also reported that the cultivars of *C. chinense* have the highest capsaicinoid levels among all *Capsicum* species (Bosland et al., 2012; Bosland & Baral, 2007; Wahyuni et al., 2011; Zhu, Sun, Cai, et al., 2019). However, the sample number and genetic diversity in previous studies were relatively low (Bosland et al., 2012; Bosland & Baral, 2007; Tanaka et al., 2017). In addition, most previous studies have utilized the whole fruit or placenta to evaluate the capsaicinoid content; thus, the contribution of the pericarp to pepper pungency was hard to evaluate (Bosland et al., 2012; Canto-Flick et al., 2008; Duelund & Mouritsen, 2017). In this study, 32 accessions from five *Capsicum* species were systemically assessed, and capsaicinoid levels in the placenta, pericarp and whole fruit were measured separately. Notably, our results show that most *C. chinense* accession pericarps contain a high level of capsaicinoids, which greatly contributes to the extreme pungency of this species. Previous studies found that a large amount of capsaicinoids accumulated on the pericarp of the cultivars ‘Trinidad Moruga’, ‘Trinidad Moruga Scorpion’ and ‘Trinidad Moruga Scorpion Yellow’ (Bosland et al., 2015; Tanaka et al., 2017). By comparing the morphological structures of the fruit pericarp and the placental tissue of extremely high-pungency cultivars (‘Trinidad Moruga’ and ‘Trinidad Moruga Scorpion’) to those of low-pungency cultivars (Jalapeno and bell pepper), the results showed that these ‘superhot’ peppers have developed accessorial vesicles on the pericarp tissue in addition to the vesicles on the placental tissue, thus leading to exceedingly high pungency in these plants (Bosland et al., 2015). Microscopic analyses have demonstrated that the elongated epidermal cells on the internal surface of the pericarp may secrete capsaicinoids (Sugiyama, 2017). However, some studies have also demonstrated that even in *C. chinense* species, some pepper cultivars have low capsaicinoid content or even nonpungency, and this mainly arises from the variation or mutation of capsaicinoid biosynthesis genes (such as *AMT* and *AT3*) (Stewart et al., 2007; Tanaka et al., 2019). Nevertheless, an understanding of the capsaicinoid content accumulation patterns in fruits of different genotypes, especially of the capsaicinoid content accumulation in the pericarp, could provide novel insight into capsaicinoid biosynthesis and accumulation in *Capsicum* fruits. The characterized *Capsicum* materials could provide valuable references for the genetic improvement of the pungency level in *Capsicum*.

### Evaluation of pungency levels (SHU) in 32 pepper accessions

To better compare the pungency levels in this study with previously reported results (Bosland et al., 2012; Duelund & Mouritsen, 2017), the total capsaicinoid levels were transformed into Scoville Heat Units (SHUs). The pungency levels (SHUs) of the placenta, pericarp and whole fruit of 32 accessions are presented in Fig. 3. Among the 32 accessions of *Capsicum* species, we found that the SHU values in the placenta (166,963 SHU~2,667,873 SHU) (Fig. 3A) were consistently higher than those in the pericarp (22,343 SHU~2,222,119 SHU) (Fig. 3B) and whole fruit (60,749 SHU~1,908,690 SHU) (Fig. 3C). As expected, the highest pungency levels were observed in accessions of *C. chinense,* followed by accessions of *C. pubescens*, *C. frutescens*, *C. annuum,* and *C. baccatum* (Fig. 3). Similarly, some results also found that *C. chinense* accumulates more capsaicinoids than accessions from *C. frutescens*, *C. annuum* and *C. baccatum* (Sarpras et al., 2016; Wahyuni et al., 2011). For most *C. chinense* accessions, the pungency levels of the placenta, pericarp and whole fruit were comparable (Fig. 3). In contrast, the pungency levels of the placenta were markedly higher than those of the pericarp and whole fruit in accessions from the other four *Capsicum* species.

**Fig. 3.**
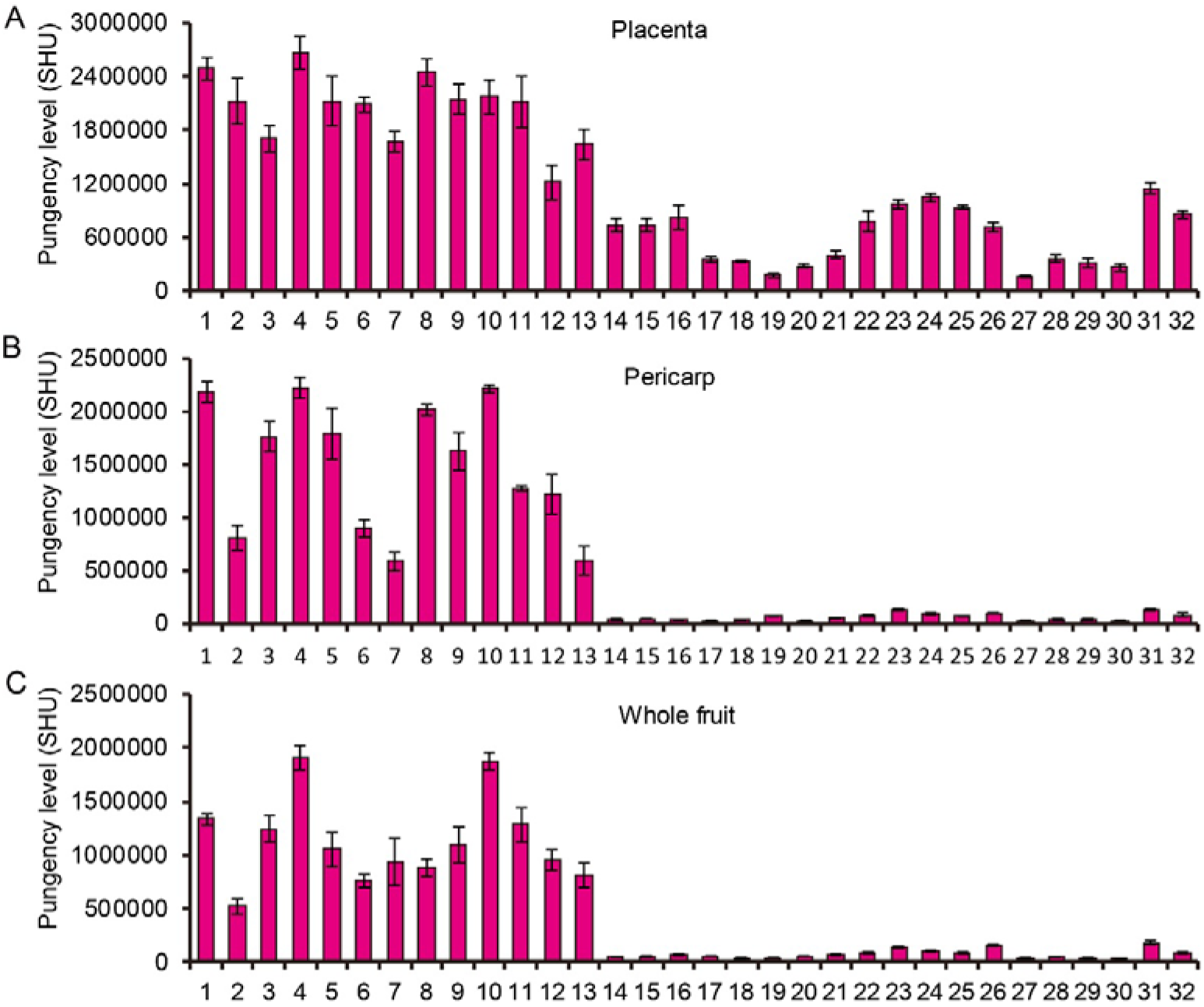
Pungency levels of 32 accessions as measured in SHUs. Pungency level in the placenta (A), pericarp (B) and whole fruit (C). Samples 1-13 are *C. chinense*, samples 14-22 are *C. annuum*, samples 23-26 are *C. frutescens*, samples 27-30 are *C. baccatum*, and samples 31 and 32 are *C. pubescens*. Data are presented as the mean ± SD (n=3).

The pungency levels of the placenta, pericarp and whole fruit of the hottest Chinese pepper, ‘Shuanla’ (sample 9), were 2,143,294 SHU, 1,623,584 SHU and 1,096,886 SHU, respectively, which are similar to the levels of the cultivar ‘Genghis Khan Brain’ (sample 5, AIASP association, Italy). In addition, ‘Bhut Jolokia’ is a group of peppers originating from India and contains many landraces ^36^, and its pungency level ranges from 520,000 SHU to 1,280,000 SHU, which is similar to previously reported results ^4, 23, 27, 37^. ‘Carolina Reaper’ is the hottest pepper, and the pungency levels of its placenta, pericarp and whole fruit reached 2,667,873 SHU, 2,222,119 SHU and 1,908,690 SHU, respectively. This pungency level is consistent with the 2,200,000 SHU reported by the Guinness Book of World Records in 2013. In Denmark, the capsaicinoid levels of ‘Carolina Reaper’ and ‘Trinidad Scorpion’ were only 1,046,000 SHU and 673,000 SHU, respectively (Duelund & Mouritsen, 2017). The significant difference in pungency values among the different regions can be explained by the fact that the environment is an important factor in determining the accumulation of capsaicinoids. Specifically, the growth conditions of Denmark are quite different from those of other regions; Denmark has a temperate climate, and the temperature for pepper growth is lower (Duelund & Mouritsen, 2017). In this context, growth conditions are essential in the measurement of capsaicinoid content and the discovery of the hottest peppers in the world. Therefore, a systematic re-evaluation of pungency values will provide robust evidence to support the pungency rankings for peppers.

### Activation of *MYB31* in the *C. chinense* pericarp elevated the expression of CBGs

The capsaicinoid biosynthesis pathway is almost completely understood and is composed of the phenylpropanoid and branched-chain fatty acid pathways (Fig. 4A). Our previous studies have demonstrated that *MYB31* is the master regulator of capsaicinoid biosynthesis and that natural variations in the *MYB31* promoter increase its expression in the *C. chinense* placenta, which finally determines the evolution of extremely pungent peppers (Zhu, Sun, Cai, et al., 2019). Given that accessions of *C. chinense* contain higher capsaicinoid levels in both the pericarp and placenta than accessions from the other four species, we hypothesized that the extreme pungency of the *C. chinense* pericarp may be attributed to the upregulation of the master regulator *MYB31* and of CBGs (Aluru et al., 2003; Arce-Rodríguez & Ochoa-Alejo, 2017; Tanaka et al., 2017; Zhu, Sun, Cai, et al., 2019). To this end, the *MYB31* and CBG transcription levels in 16 DPA fruit pericarps and placentas of the 32 accessions were analyzed. The results indicated that the expression levels of *MYB31* and CBGs varied greatly in the placentas of the 32 accessions, and the expression was consistently higher among accessions of *C. chinense* (Fig. 4B). This finding is in agreement with the highest CAP, DHCAP and total capsaicinoid contents found in this species. Notably, in the pericarp of the *C. chinense* accessions, the expression levels of key CBGs (such as *AMT*, *BCAT*, and *AT3*) and *MYB31* were remarkably upregulated (Fig. 4C). To further confirm that the transcription levels of *MYB31* and CBGs were associated with the capsaicinoid content, a correlative analysis was performed. The results indicated that *MYB31* and CBG transcription levels in the placenta were positively correlated with the total capsaicinoid content; the correlation coefficient (*R*^2^) ranged from 0.65 to 0.82 (Fig. 4D). In addition, the *MYB31* and CBG transcription levels were significantly correlated with the total capsaicinoid contents in the pericarp, and the correlation coefficients (*R*^2^) ranged from 0.55 to 0.85 (Fig. 4E). Moreover, we found that *MYB31* and CBG transcript abundance in both the placenta and pericarp exhibited intraspecific as well as interspecific variations that were consistent with the capsaicinoid content levels. It is generally recognized that capsaicinoids are produced exclusively in the placenta of the fruit; thus, CBGs are highly or specifically expressed in the fruit of the placenta from 16 DPA to the mature green stage (Fujiwake et al., 1980; Kim et al., 2014; Qin et al., 2014; Suzuki et al., 1980; Zhu, Sun, Cai, et al., 2019). However, in this study, almost all key CBGs (*AMT*, *KasI* and *AT3*) were upregulated in the pericarp of the *C. chinense* accessions but weakly expressed in the pericarp of the other four *Capsicum* species (Fig. 4C). Although published research has shown that CBGs are expressed in the cultivar ‘Trinidad Moruga Scorpion Yellow’ pericarp (Tanaka et al., 2017), the upregulation of CBGs in the pericarps of other cultivars and the role of MYB31 in this process are still far from understood. More importantly, we found that capsaicinoid levels were controlled at the transcriptional level and were consistent with master regulator *MYB31* expression, which is also upregulated in the pericarp of *C. chinense* accessions (Fig. 4C). In the pericarp, the correlation coefficient of the *MYB31* transcription level and the total capsaicinoid content reached *R*^*2*^ = 0.85, strongly supporting the hypothesis that *MYB31* plays an important role in regulating the CBGs to control the biosynthesis of capsaicinoids in the pericarp. Although the capsaicinoid content in *C. chinense* accessions was higher than that in the accessions of the other four *Capsicum* species, we found that the expression of some CBGs (such as *BCKDH* and *FatA*) in some *C. chinense* accessions was lower than that in some accessions of other species. A possible explanation is that the capsaicinoid content is a quantitative trait^3, 5, 36^ that is controlled by multiple QTLs and/or genes. Nevertheless, understanding the underlying molecular mechanisms behind capsaicinoid content variation will provide an effective strategy for increasing the capsaicinoid content in pepper breeding.

**Fig. 4.**
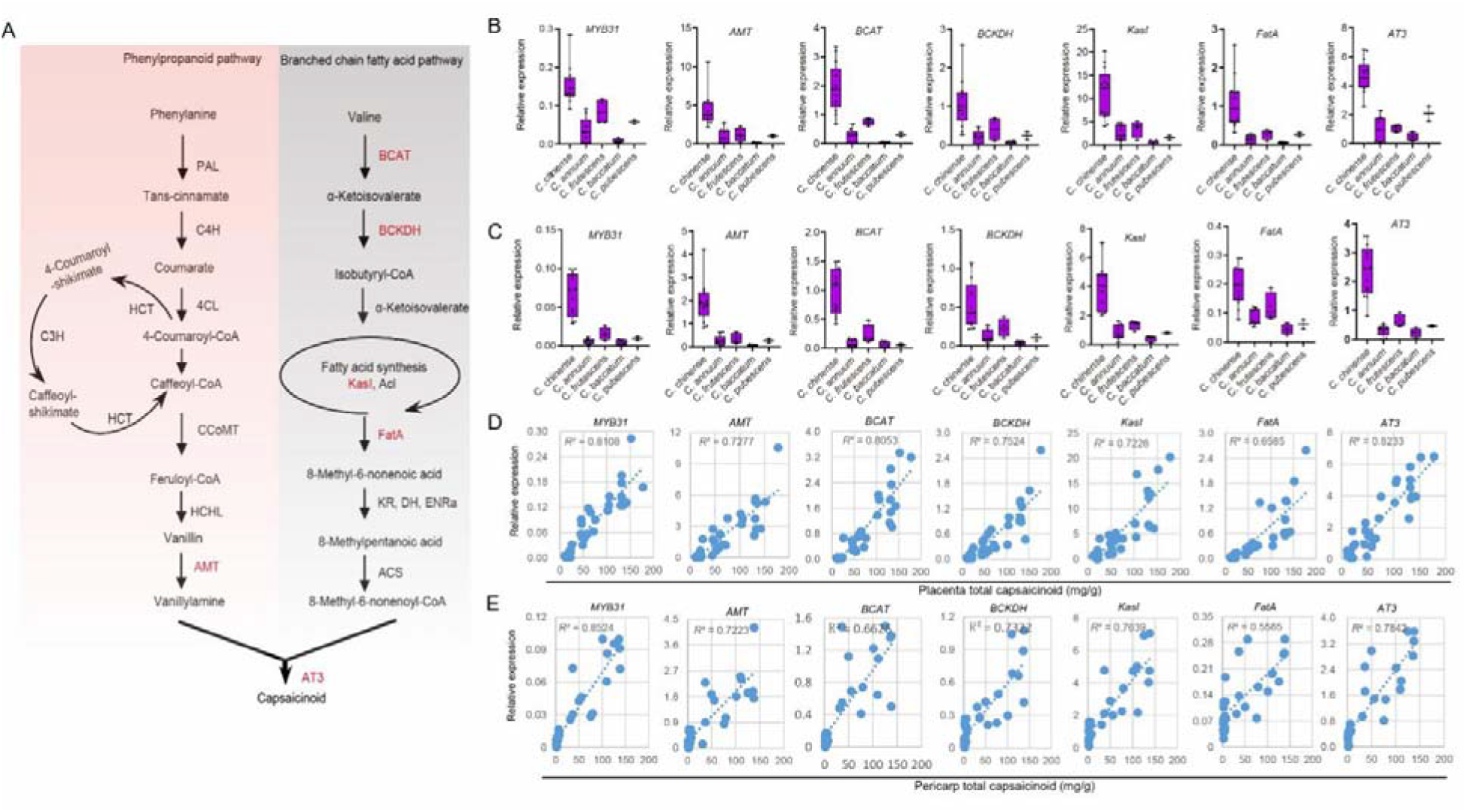
Transcriptional analysis of the expression of CBGs and the master regulator *MYB31*. (A) Schematic depiction of the capsaicinoid biosynthetic pathway. PAL, Phe ammonia-lyase; C4H, cinnamate 4-hydroxylase; 4CL, 4-coumaroyl-CoA ligase; HCT, hydroxycinnamoyl transferase; CoMT, caffeic acid 3-O-methyltransferase; HCHL, hydroxycinnamoyl-CoA hydratase lyase; AMT, aminotransferase; BCAT, branched-chain amino acid aminotransferase; BCKDH, branched-chain α-ketoacid dehydrogenase; KasI, ketoacyl-ACP synthase I; Acl, acyl carrier protein; FatA, acyl-ACP thioesterase; ENRa, enoyl-ACP reductase a; KR, ketoacyl-ACP reductase; DH, hydroxyacyl-ACP dehydratase; ACS, acyl-CoA synthetase; AT3, acyltransferase 3. (B) Transcriptional analysis of CBGs and the master regulator *MYB31* in the 16 DPA fruit placenta of *C. chinense*, *C. annuum*, *C. frutescens*, *C. baccatum* and *C. pubescens* samples. (C) Transcriptional analysis of CBGs and the master regulator *MYB31* in the 16 DPA fruit pericarp of *C. chinense*, C. annuum, *C. frutescens*, *C. baccatum* and *C. pubescens* samples. (D) The correlation of the CBGs and the *MYB31* transcription level with the total capsaicinoid contents in the placenta. (E) The correlation of the CBGs and the *MYB31* transcription level with the total capsaicinoid contents in the pericarp.

### Capsaicinoid accumulation patterns and *MYB31* and CBG expression in the placenta and pericarp of two genotypes during different developmental stages

To provide further insights into the transcriptional regulation of capsaicinoid biosynthesis in the pericarp, the capsaicinoid accumulation patterns and gene expression levels in different developmental stages of two genotypes of peppers were investigated. *C. annuum* and *C. chinense* are considered the most widely cultivated pepper species in the world (Zhu, Sun, Wei, et al., 2019). The capsaicinoid contents in fruits in different developmental stages of the ‘59’ inbred line (Sample 14) and ‘Trinidad Moruga Scorpion’ (Sample 1) were analyzed (Fig. 5). The placenta and pericarp were separated from the pepper fruits at different developmental stages and sampled for capsaicinoid content detection. In both ‘Trinidad Moruga Scorpion’ and the ‘59’ inbred line, the CAP, DHCAP and total capsaicinoids in the placenta started to accumulate at 10 DPA and 16 DPA, reached a peak at 45 DPA and then slightly decreased at 55 PDA (Fig. 5C and 5E). In the pericarp, peaks in CAP and DHCAP contents were observed at 45 DPA in both genotypes (Fig. 5D and 5F). Our findings are consistent with results reported previously(Barbero et al., 2014; Fayos et al., 2019). In the late developmental stage of fruits, capsaicinoids in the ‘Trinidad Moruga Scorpion’ pericarp were extremely abundant (Fig. 5D). In contrast, in the whole developmental stage of the fruit of the ‘59’ inbred line, capsaicinoids were almost undetectable in the pericarp (Fig. 5F). In particular, in the pericarp of the 45 DPA fruit, the ‘Trinidad Moruga Scorpion’ capsaicinoid level was 60-fold higher than the ‘59’ inbred line level, while it was only 3-fold higher between the two placentas (Fig. 5C-5F). During different fruit developmental stages, the capsaicinoid contents in the placenta and pericarp were at relatively comparable levels in ‘Trinidad Moruga Scorpion’ (Fig. 5C and 5D). In contrast, the capsaicinoid levels in the placenta of the ‘59’ inbred line during different fruit developmental stages were significantly higher (approximately 20-fold) than those in the pericarp (Fig. 5E and 5F).

**Fig. 5.**
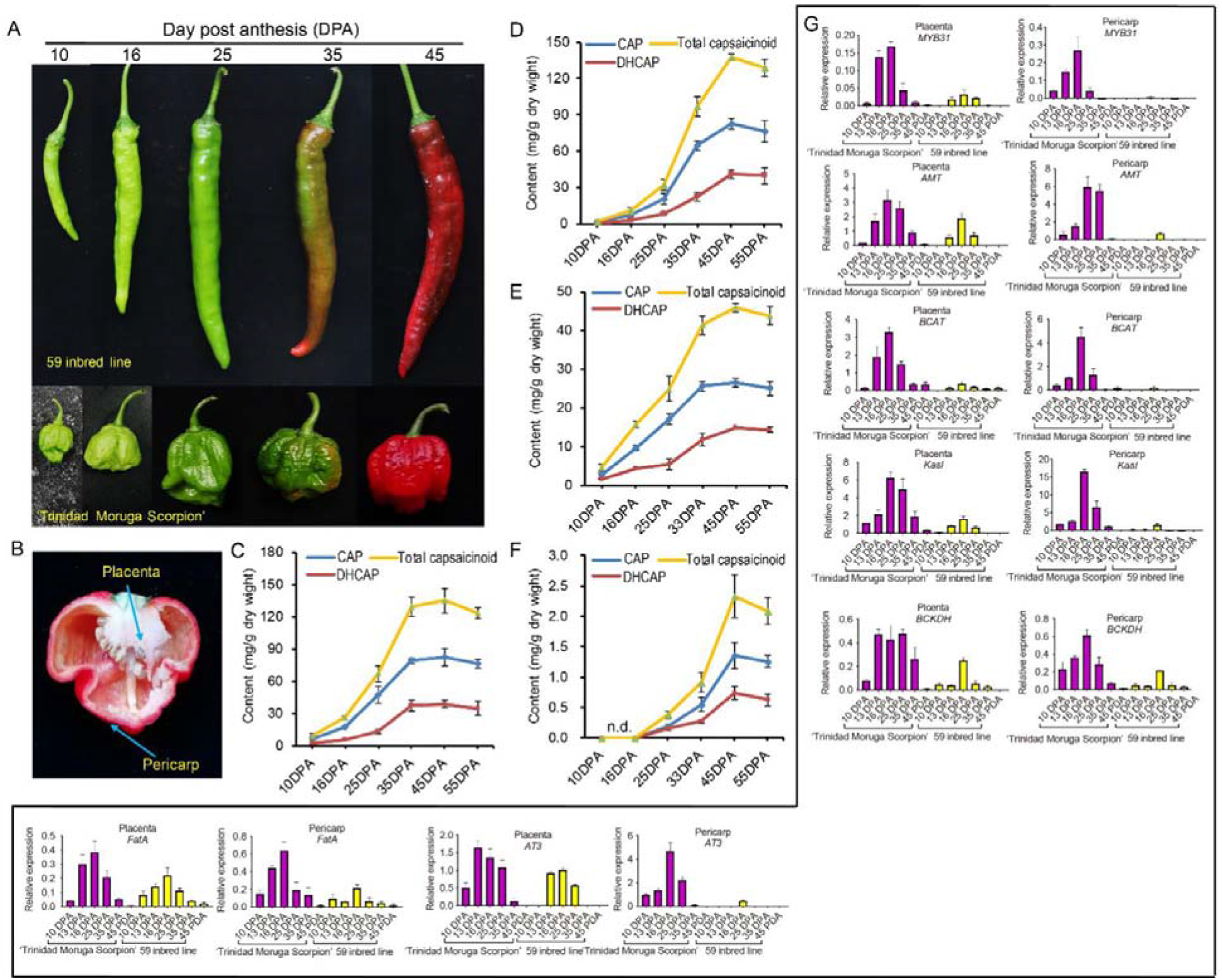
Capsaicinoid content in the pepper placenta and pericarp at different developmental stages. (A) Five developmental stages of ‘Trinidad Moruga Scorpion’ and ‘59’ inbred line fruits. (B) Cross-section of ‘Trinidad Moruga Scorpion’ fruit at 45 DPA. (C) and (D) CAP, DHCAP and total capsaicinoid contents in the ‘Trinidad Moruga Scorpion’ placenta (C) and pericarp (D). (D) and (F) CAP, DHCAP and total capsaicinoid contents in the ‘59’ inbred line fruit placenta (E) and pericarp (F). Data from (B-F) are presented as the mean ± SD (n=3). n.d., undetected. (G) Transcriptional analysis of CBGs and the master regulator *MYB31* in the placenta and pericarp of ‘Trinidad Moruga Scorpion’ and the ‘59’ inbred line at different developmental stages. Data are presented as the mean ± SD (n=3).

To address the relationship between gene expression and capsaicinoid content, qRT-PCR was used to analyze the master regulator *MYB31* and key CBG expression patterns in the placenta and pericarp. The results revealed similar expression patterns of CBGs between ‘Trinidad Moruga Scorpion’ and ‘59’ inbred line placenta tissues, and the CBGs were primarily expressed from 13 DPA to 35 DPA (Fig. 5G). The expression level of CBGs in ‘Trinidad Moruga Scorpion’ placenta was higher than that in the ‘59’ inbred line (Fig. 5B), which is consistent with the higher capsaicinoid levels in ‘Trinidad Moruga Scorpion’ (Fig. 5C and 5D). In addition, the master regulator *MYB31* was also significantly upregulated in the ‘Trinidad Moruga Scorpion’ pericarp and placenta (Fig. 5G). To date, it is generally recognized that CBGs are highly or specifically expressed in the placenta and rarely expressed in other tissues (Aluru et al., 2003; Fujiwake et al., 1980; Kim et al., 2014; Stewart et al., 2005; Suzuki et al., 1980). Notably, we found that the master regulators *MYB31* and CBGs are highly expressed in ‘Trinidad Moruga Scorpion’ fruit pericarps at 13 DPA-25 DPA, which is consistent with the capsaicinoid accumulation during this stage. However, the transcription levels of *MYB31* and key CBGs were low in the ‘59’ inbred lines at different developmental stages of the fruit pericarp (Fig. 5G). Previously, studies found that CBGs are highly or specifically expressed in the placenta from 16 DPA to the mature green stage, and capsaicinoids are produced exclusively in glands on the placenta of the fruit (Aluru et al., 2003; Fujiwake et al., 1980; Iwai et al., 1979; Stewart et al., 2007; Suzuki et al., 1980). Some fatty acid synthase genes, such as *Acl*, *FatA* and *Kas*, were positively correlated with the degree of pungency (Aluru et al., 2003). Recently, a comparison between *C. chinense* (‘Bhut Jolokia’), *C. frutescens* and *C. annuum* suggested that higher levels of CBG expression might be responsible for the extremely high pungency (Sarpras et al., 2016). Another study investigated ‘Trinidad Moruga Scorpion Yellow’ pungency levels and found that its pericarp exhibited a markedly higher capsaicinoid concentration than those of the other three cultivars ^32^. Expression analysis revealed that CBGs were expressed exclusively in the ‘Trinidad Moruga Scorpion Yellow’ pericarp. Our results indicate that the upregulation of CBG expression and the elevation of capsaicinoid levels in the ‘Trinidad Moruga Scorpion’ pericarp arise from the activated expression of the master regulator MYB31, which leads to the CBGs being strongly activated in the pericarp and increases the metabolic production of capsaicinoid, which results in the extreme pungency of the pericarp.

## Conclusions

In this study assessing 32 accessions, the high pungency level in the accessions of *C. chinense* is attributed to the high levels of capsaicinoids accumulating in both the placenta and the pericarp; in particular, the capsaicinoids derived from the pericarp make a major contribution to the pungency of *C. chinense*. The transcription levels of *MYB31* and CBGs were positively correlated with capsaicinoid levels. The high capsaicinoid content in the *C. chinense* pericarp is attributed to the upregulation of the master regulator *MYB31*, which leads to the strong activation of CBG expression. The capsaicinoid accumulation trends in ‘Trinidad Moruga Scorpion’ and ‘59’ in different developmental stages of the placenta and pericarp were consistent with the *MYB31* and CBG gene expression levels. Taken together, our study not only provides new insights into the capsaicinoid accumulation patterns in pepper fruits of different genotypes but also lays a foundation for the genetic manipulation of pepper capsaicinoid content for various purposes.

## Acknowledgments

This work was supported by the National Key Research and Development Program (2018YFD1000800) and the National Natural Science Foundation of China (31572124). In addition, Zhangsheng Zhu wants to acknowledge the patience, care and support received from Binmei Sun over the years.

Appendix A. Supplementary data

---

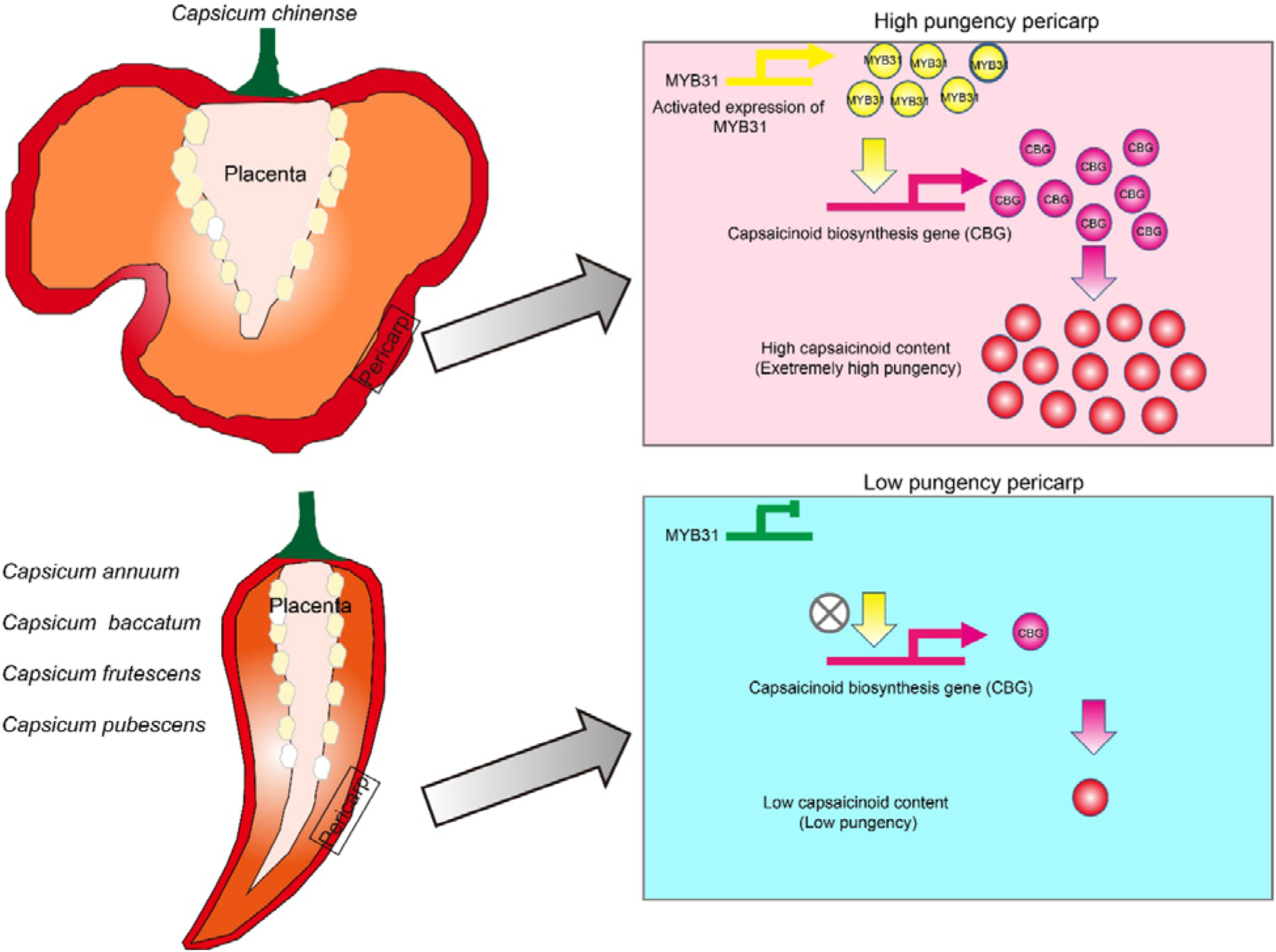

